# LINCS gene expression signatures analysis revealed bosutinib as a radiosensitizer of breast cancer cells by targeting *eIF4G1*

**DOI:** 10.1101/2020.04.24.059378

**Authors:** Sai Hu, Pingkun Zhou, Xiaodan Liu, Xiaoyao Yin, Dafei Xie, Bo Huang, Hua Guan

## Abstract

Radioresistance represents the predominant cause for radiotherapy failure and disease progression, resulting in increased breast cancer mortality. Through gene expression signatures analyses of Library of Integrated Network-Based Cellular Signatures (LINCS) and Gene Expression Omnibus (GEO), the present study aimed to identify potential candidate radiosensitizers from known drugs systematically. The similarity of integrated gene expression signatures between irradiated *eIF4G1*-silenced breast cancer cells and known drugs was measured by enrichment scores. Drugs with positive enrichment scores were selected as potential radiosensitizers. The radiosensitizing effects of the candidate radiosensitizers were analyzed in breast cancer cells (MCF-7, MX-1, and MDA-MB-231) by CCK-8 assays and colony-forming capability after exposure to ionizing radiation. Cell apoptosis was detected by flow cytometry. Expressions of *eIF4G1* and a series of DNA damage response proteins were analyzed by Western blot assays. Bosutinib was proposed to be a promising radiosensitizer as its administration markedly reduced the dosages of both the drug and ionizing radiation and was associated with fewer adverse drug reactions. The combined treatment with ionizing radiation and bosutinib significantly increased the cells killing potency in all three cell lines as compared to ionizing radiation or bosutinib alone. MX-1 cells were revealed to be the most sensitive to both ionizing radiation and bosutinib among the three cell lines. Bosutinib noticeably downregulated the expression of *eIF4G1* in a dose-dependent manner and also reduced the expression of DNA damage response proteins (including *ATM*, *XRCC4*, *ATRIP*, and *GADD45a*). Moreover, *eIF4G1* could be a key target of bosutinib through which it regulates DNA damage induced by ionizing radiation. Thus, taken together, bosutinib may serve as a potential candidate radiosensitizer for breast cancer therapy.

## Introduction

Currently, breast-conservative surgery, followed by radiation therapy (RT), represents the standard of care for breast cancer[1]. Although breast-conservative surgery with RT improves the overall survival, efficacy and safety of RT remain controversial and unfavorable, mainly due to the survival of cancer cells that develop resistance to RT[1–4]. Moreover, the repopulation of these radioresistant cells leads to treatment failure and local recurrence, thus, requiring administration of higher doses of radiation to repopulation regions, which may cause damage to healthy tissues surrounding the tumors. Therefore, the identification of new pharmacological approaches to overcome radioresistance of cancer cells is highly desired[3].

It is well recognized that intrinsic and ionizing radiation (IR)-induced radioresistantce determines cellular responses to RT. Previous studies have identified a series of radioresistance-associated genes, including *eIF4G1*[5], *IGF-1R*[6], *p53*[7], and *DNA-PK*, which represented potential molecular targets for radiosensitization[8]. DNA repair, cancer stem cells, apoptosis[9], cell cycle arrest[10,11], autophagy[12], and hypoxia[13,14] were indicated as crucial biological mechanisms leading to radioresistance of cancer cells. Among the radiosensitizers being actively investigated and those in use are agents targeting hypoxic cells, inhibitors of ion channels[15], regulators of proteins (such as enzymes)[4,10,16] and autophagy reducers[17]. However, difficulties exist with these compounds are their lack of specificity, elusive underlying molecular mechanisms, and undesired effects[3]. Moreover, there is a paucity of promising radiosensitizers for a variety of cancer types, including breast cancer.

The prevailing paradigm for drug discovery, including radiosensitizers, is extremely time-consuming and expensive; besides, the design and synthesis of new compounds, along with high-throughput screening, remain challenging [18]. Drug repositioning (also known as drug repurposing or drug reprofiling) is intended to predict alternative implications for a pioneering drug. This strategy would help reduce the time and cost associated with new drug development and advance the delivery of new drugs to patients with severe diseases[19]. Thus, drug repositioning is a promising strategy to identify and develop known compounds as potentially effective radiosensitizers to improve the efficacy and reliability of RT.

In the present study, we proposed bosutinib, a small molecular *BCR-ABL* kinase inhibitor approved by the US Food and Drug Administration (FDA) in 2012, as a potential radiosensitizer of breast cancer for eliciting similar therapeutic effects as silencing of *eIF4G1* following IR and has been demonstrated to significantly sensitize cancer cells to DNA damage induced by RT through regulation of cell apoptosis[20], and expression of proteins including *eIF4G1*, *ATM*, and *XRCC4*. In this study, we conducted a series of mechanistic experiments and verified the synergistic effect of bosutinib and γ-ray on breast cancer cells. The combination of these two showed an apparent anti-tumor effect in breast cancer cells at relatively lower effective dosages of both bosutinib and radiation with fewer side effects.

## Materials and methods

### The Library of Integrated Network-based Cellular Signatures (LINCS)

Cellular signatures of all compounds used in this study were collected and extracted from a comprehensive systems biology program National Institutes of Health (NIH) Library of Integrated Network-based Cellular Signatures (LINCS) (https://commonfund.nih.gov/LINCS). The LINCS datasets comprise cellular response signatures of 22412 unique perturbations applied to 56 different cellular contexts, among which 16425 are chemical reagents, including drugs, ligands, and other small molecules applied at different time points and doses[21]. The L1000 technology, a high-throughput method, allows estimating genome-wide mRNA expression, and these L1000 signatures may facilitate the development of computational models for complex diseases and drug effects.

### Gene Expression Omnibus (GEO)

The gene expression profile dataset GSE41627 used in this study was extracted from the Gene Expression Omnibus (GEO) (https://www.ncbi.nlm.nih.gov/geo/), an international public functional data repository for submission, storage, and integration of high-throughput gene expression data from gene expression and genomic hybridization experiments. GEO comprises three domains, including platforms, series, and samples. A total of 20816platforms, 128393 series, and 3548760 samples in 4348 datasets are available currently[22].

### Calculation of enrichment scores (ES)

Both gene IDs in GSE41627 and LINCS were converted into an official gene symbol according to different platforms. Enrichment scores (ES) calculated by *R* code indicated the similarity between gene expression signatures of cells treated with a variety of compounds and irradiated *eIF4G1*-silenced cells. Default parameters in gene set enrichment analysis desktop application[23] were used to generate the gene rank list for each profile. The classic scoring scheme was applied to calculate ES for the silencing of *eIF4G1* and compounds in LINCS. The significance of ES was assessed with 1000 permutations of the data.

### Cell culture and irradiation

The human breast cancer cell lines, including MX-1, MCF-7, and MDA-MB-231, were used in this study. MCF-7 and MDA-MB-231 cells were purchased from the American Type Culture Collection (Manassas, VA, USA). MX-1 cells were provided by the Beijing Institute of Transfusion Medicine. The cells were cultured in Dulbecco’s modified eagle medium (DMEM) supplemented with 10% fetal bovine serum (Sigma), 100U/mL penicillin and 100μg/L gentamycin and incubated at 37℃ in a humidified atmosphere containing 5% CO_2_. Cells were divided into four groups: irradiation alone, drug alone, irradiation and drug, and the control groups. The cells of irradiation groups were irradiated with ^60^Co γ-rays at a dose rate of 1.98 Gy/min at room temperature in the Beijing Institute of Radiation Medicine.

### Drugs and antibodies

Drugs used in this study were as follows: bosutinib (E047103, EFEBIO Shanghai, China), bifonazole (B3563, Sigma, USA), and isosorbide (652-67-5, J&K, Beijing, China). Antibodies used were as follows: anti-*eIF4G1* (ab2609, Abcam, Cambridge, UK), anti-*eIF4G2* (P78344, Cell Signalling Technology, USA), anti-*Chk1* (2360s, Cell Signalling Technology), anti-phospho-*Chk1* (S345; 2341s, Cell Signalling Technology), anti-*Chk2* (2662s, Cell Signalling Technology), anti-phospho-*Chk2* (Thr68; 2197s, Cell Signalling Technology), anti-*H2AX* (S139; ab11174, Abcam, UK), and anti-*β-actin* (TA-09, Beijing Zhongshan Jinqiao Biotechnology Co. Ltd.).

### Cell proliferation and survival analyses

Cell proliferation was analyzed using a Cell-Counting Kit-8 assay kit (CCK-8; Engreen, China) following the manufacturer’s instructions. Briefly, cells were seeded in 96-well plates and exposed to increasing concentrations of tested drugs alone. To assess cell proliferation, CCK-8 solution (CK04, DOJINDO, Japan) was added at 48 h and 72 h after the treatment, respectively. The absorbance was measured at 450 nm using a microplate reader. Half-maximal inhibitory concentrations (IC50) were calculated by Graphpad prism software (Systat Software, Inc.). Each experiment was performed in triplicate.

Cell survivals were assessed by the colony formation assays. Cells were seeded into 60-mm culture plates at the density of 3×10^2^ cells/plate and exposed to various concentrations of bosutinib for 24 h. After irradiation, the cells were subsequently cultured in normal medium for 10-15 days. Surviving tumor cells were fixed with ethanol and stained with Giemsa and then counted.

### Immunoblotting analyses

Total proteins were extracted from the cultured cells on ice with lysis buffer supplemented with protease inhibitors cocktail. Cell lysates were separated using sodium dodecyl sulfate-polyacrylamide gel electrophoresis (SDS-PAGE) with a Bio-Rad Bis-Tris Gel system and then transferred the proteins to polyvinylidene difluoride membrane. Separated proteins were then transferred to polyvinylidene fluoride (PVDF) membranes. The membranes were blocked with 5% non-fat dry milk at room temperature for 1h. Subsequently, the membranes were incubated with following rabbit monoclonal primary antibodies against *eIF4G1*, *eIF4G2*, and mouse monoclonal antibody against *β-actin* at 1:3000, 1:1000, and 1:1000 dilution, respectively. Then, the membranes were washed three times with TBST and incubated with indicated secondary antibodies conjugated with horseradish peroxidase (HRP) (1:4000, 1h) at room temperature. The target bands were visualized using the Image Quant LAS500 system.

### Cell apoptosis analyses

Cells (3×10^5^−6×10^5^) were treated with bosutinib for 24 h before irradiation with 4 Gy ^60^Co γ-rays and then subjected to trypsinization with trypsin/EDTA (Hyclone) at 8, 12, and 24 h after irradiation, respectively. For apoptosis analyses, the cell pellets were re-suspended in 100 μl of binding buffer and stained with 5 μl of PI (50 μg/ml) and 5 μl of Annexin V-FITC (DA10, Dojindo, Shanghai, China). Apoptosis was then analyzed with a flow cytometer (ACEA Novocyte).

### Statistical analyses

All the experiments were performed in triplicate. Data from three or more independent experiments were expressed as the mean ± standard deviation of the mean. Data obtained from each group were tested by one-way analysis of variance (ANOVA). Differences between two groups were determined using the Student’s *t*-test. *P* < 0.05 was considered statistically significant. Statistical analyses were performed using SPSS 18.0 software (IBM, Armonk, NY, USA).

## Results

### Extraction and integration of gene expression profiles

After a thorough screening and manual curation, we selected the gene expression profiles from the GSE41627 dataset for the identification of potential radiosensitizers. This dataset reported by Badura *et al.* was generated from breast cancer cells MDA-MB-231 and MCF-7 exposed to 10 Gy of γ-ray irradiation, with or without silencing of translation initiation factor *eIF4G1*[20]. Treatment groups were divided into four groups, including irradiated cells, *eIF4G1*-silenced cells, irradiated *eIF4G1*-silenced cells, and the control groups. The silencing of *eIF4G1* has been shown to significantly sensitize breast cancer cells to IR. Therefore, radiosensitizer candidates were identified by comparison of similarity in biological effects between the gene expression profiles from the GSE41627 dataset and those of breast cancer cells from LINCS in response to treatments with different kinds of chemical compounds. The flow diagram is illustrated in Fig 1.

**Fig 1.**
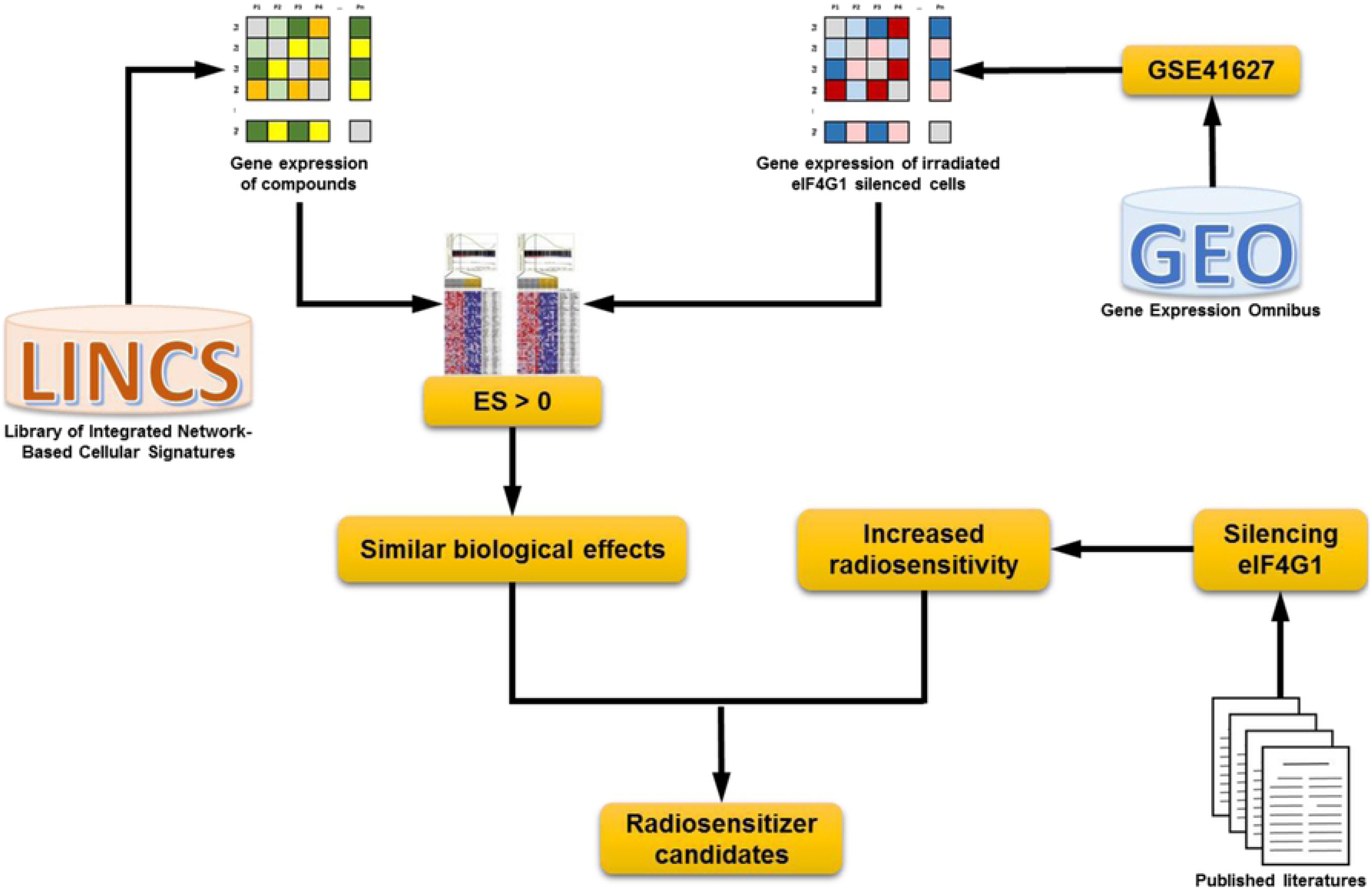
The schematic representation of the workflow of the approach in identification of candidate radiosensitizers. They were screened out from the Library of Integrated Network-based Cellular Signatures (LINCS) based on comparison of similarity of gene expression profiles between drug treated and irradiated eIF4G1-silenced breast cancer cells. Note: ES refers to enrichment score.

### Identification of potential radiosensitizers

Gene expression signatures of the GSE41627 dataset, as well as those generated from breast cancer cells treated with various compounds, were analyzed and calculated by *R* code. The similarity in biological effects between a compound and the *eIF4G1*-silenced cells after irradiation was measured by ES. A positive ES of a known compound indicates that the drug-perturbed profile is similar to the effect of silencing *eIF4G1* on cellular response to irradiation, implying that queried compounds can increase the radiosensitivity of breast cancer cells.

A total of 2089 entries including different concentrations of one drug from LINCS were evaluated. These were realized at 96 h and 144 h after irradiation. We calculated the maximum and average value of ES for each entry. Thresholds for screening were selected based on ES of each compound to ensure that no more than 5% of compounds were screened out.

In compounds at 96 h after IR exposure, both maximum and average ES of 369 drugs were positive (S1 Table). Considering 0.70 and 0.56, respectively, as thresholds for the maximum ES and average ES, two kinds of experimental drugs SU-11652 and latrunculin-b were selected. SU-11652 is a multi-targeting receptor tyrosine kinase inhibitor being investigated as an anticancer drug[24]. Latrunculin-b is a cell-permeable actin polymerization inhibitor against cancer[25]. Considering only the drugs approved by FDA, four drugs, including floxuridine, palbociclib isethionate, raltitrexed, and bosutinib were screened out when the threshold of maximum ES was set as 0.66, and the average was 0.48.

At 144 h after radiation, 179 drugs exhibited positive maximum and average ES (S1 Table). PD-0325901, an experimental *MEK* kinase inhibitor, with a marked anti-tumor activity [26] was identified with 0.49 and 0.36 as thresholds for the maximum ES and average ES, respectively. The maximum ES of 10 FDA approved drugs was above the threshold of 0.47, including lepirudin, pentoxifylline, and trametinib. While the average ES of 14 approved drugs was above threshold 0.30, including bifonazole, isosorbide mononitrate, and anastrozole.

### Selection of promising candidate radiosensitizers

The comparison of transcriptional signatures provided a wide range of candidate radiosensitizers, and Table 1 presented the preferred candidates. Compounds with both the maximum ES and average ES higher than thresholds were proposed except in the case of approved drugs at 144 h after irradiation as no compounds met the criteria. We, therefore, selected the top two compounds based on maximum ES and average ES.

**Table 1.**
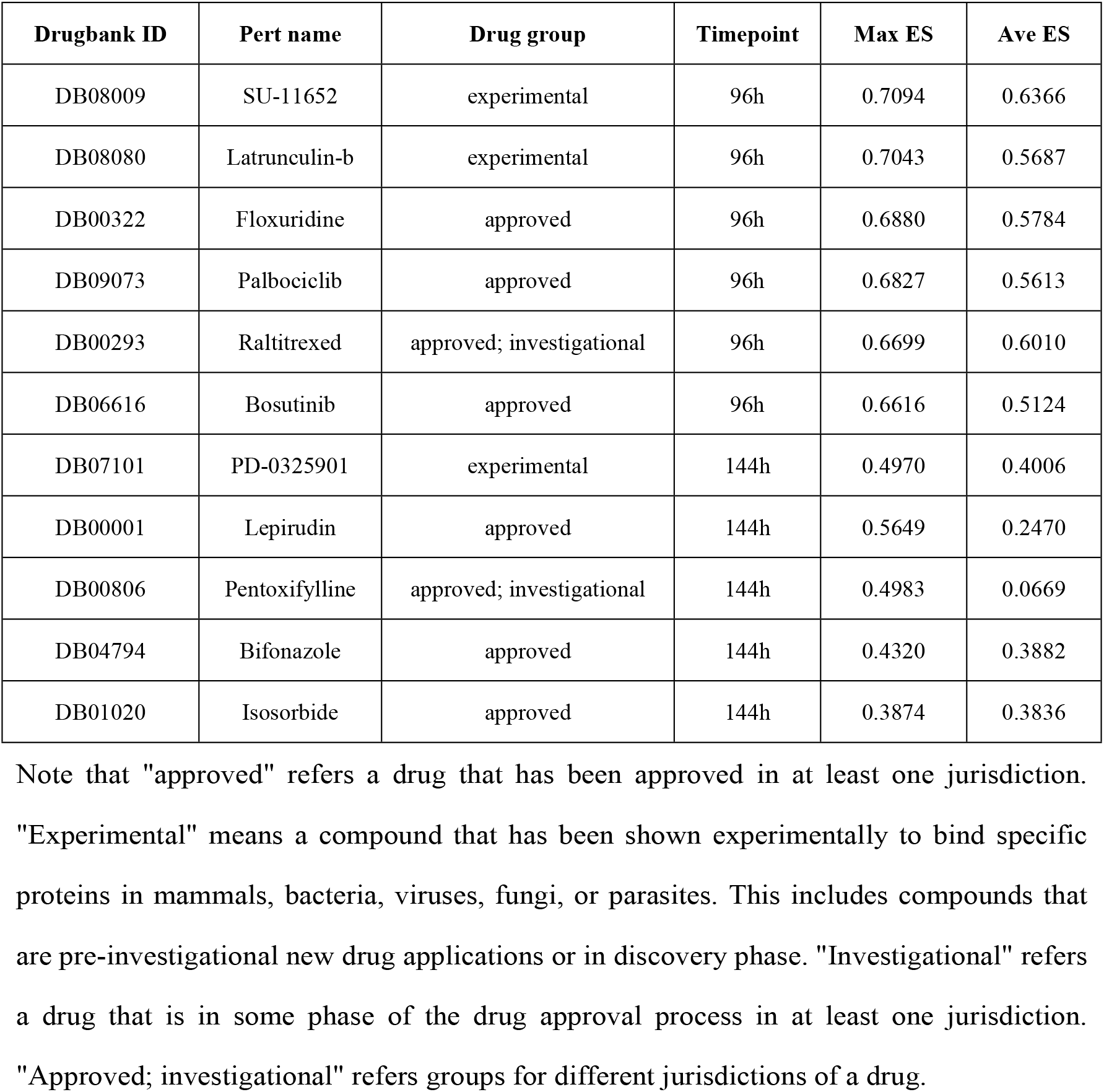
Top-ranked compounds compared with irradiated *eIF4G1*-silenced breast cancer cells.

We selected bosutinib, bifonazole, and isosorbide for further verification and additional studies on the role and mechanism of *eIF4G1* in the cellular response to IR-induced DNA damage after manual curation and decision. Among them, bosutinib has been used in the treatment of Philadelphia chromosome-positive (Ph+) chronic myelogenous leukaemia with resistance or intolerance to prior therapy, according to Drugbank. As a *BCR-ABL* kinase inhibitor, tolerance and safety of this drug have been improved compared with previous tyrosine kinase inhibitors. *BCR-ABL* kinase plays an important role in the formation and rapid progression of chronic myelogenous leukaemia and is among the primary targets of bosutinib. Other known targets include *LYN*, *SRC*, *CDK2*, and *MAP2K1.*

### Bosutinib inhibits cell growth

The cell growth inhibitory effects of the selected candidate drugs bosutinib, bifonazole, and isosorbide on MX-1 cells were presented in Fig 2. Of these, isosorbide exhibited no evident inhibitory effect at the concentration lower than 1 mM (Figs 2C and D). Bosutinib was more effective (Figs 2A and B) as its IC50 was only 1% of that of bifonazole (Fig 2D), implying for further evaluation and examination of bosutinib.

**Fig 2.**
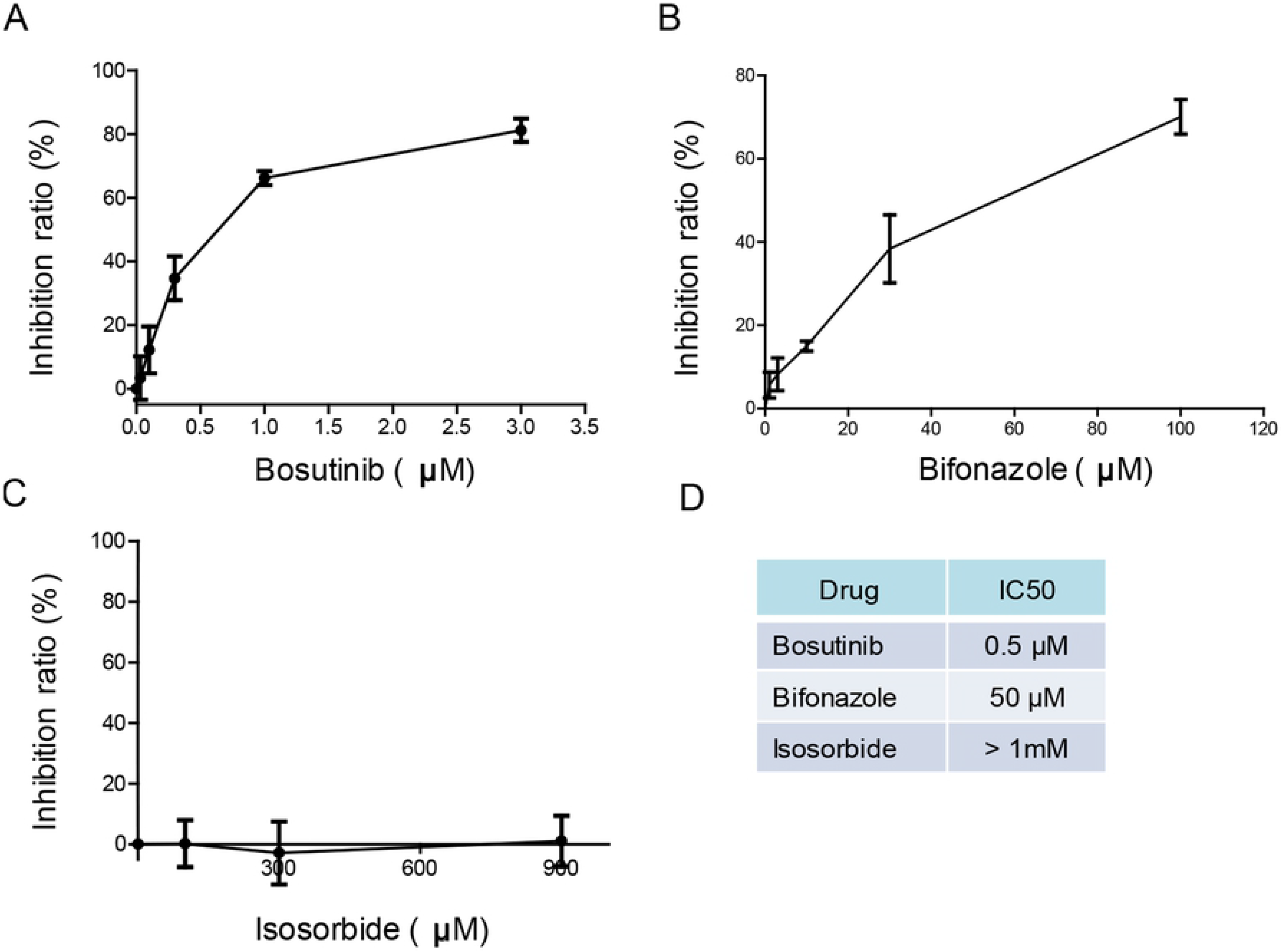
Cell growth inhibition and half-maximal inhibition concentration (IC50) of the selected candidate drugs. MX-1 cells were treated with different concentrations of bosutinib, bifonazole, or isosorbide for 48 h. The cell growth inhibition was determined by CCK-8 assays. The DMSO solvent was used as the control. (A-C) Inhibition of MX-1 cells growth by bosutinib, bifonazole, and isosorbide. (D) The IC50 of the tested drugs on cell growth. Each experiment was repeated three times.

Bosutinib was applied to two additional cell lines MCF-7 and MDA-MB-231, besides MX-1 for 48 h and 72 h, respectively. Inhibition rates altered with respect to concentrations of bosutinib, which were determined based on responses of different cells to this compound. In MX-1 and MDA-MB-231 cells, the inhibition rates increased noticeably to the concentrations of bosutinib (Figs 3A and C), while the response in MCF-7 cells was indistinct (Fig 3B). MX-1 cells were much more sensitive to bosutinib than the other two cell lines at the concentrations of 0.5μM and 1.0μM (Fig 3D). Colony formation assays were conducted in MDA-MB-231 and MCF-7 cells. It was shown that cell growth in the two cells was inhibited when the concentration of bosutinib increased (Figs 3E and F). MDA-MB-231 cells were revealed to be more sensitive to bosutinib than MCF-7 cells (Fig 3F).

**Fig 3.**
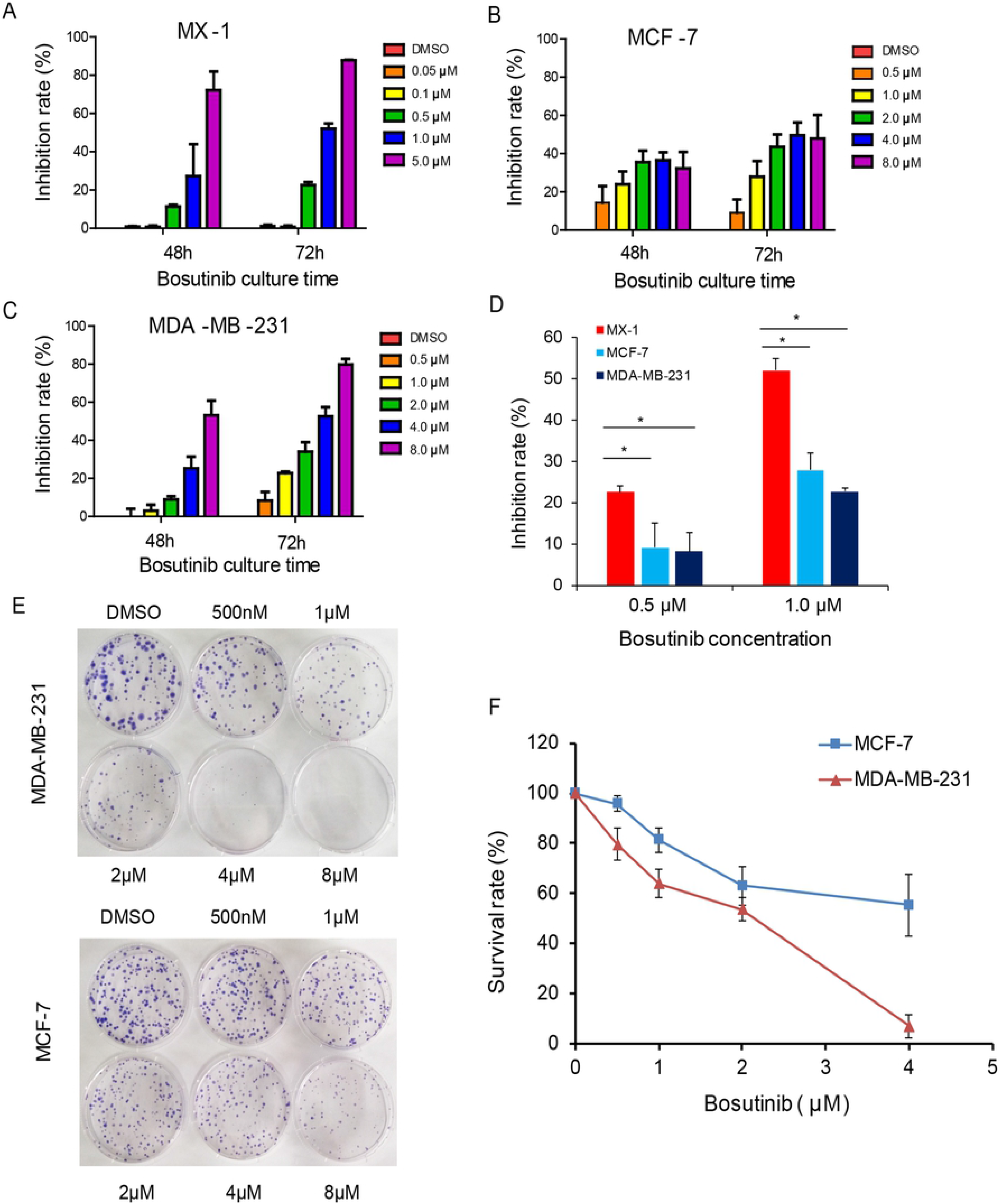
Effects of bosutinib on the cell growth of different cell lines. (A-C) Bosutinib inhibits the cell growth of MX-1, MCF-7, and MDA-MB-231, determined by CCK-8 assays. (D) Comparison of cell growth inhibition by bosutinib on the three tested cell lines. (E) Effect of bosutinib on colony formation of MCF-7 and MDA-MB-231 cells. (F) Cell survival of MCF-7 and MDA-MB-231 cells treated with different concentrations of bosutinib represented by cell colony-forming capability. \**P* < 0.01.

### Bosutinib increases radiosensitivity of breast cancer cells

Clonogenic assays were used to investigate the radiosensitivity of MCF-7 and MDA-MB-231 cells. The cells were treated with bosutinib at a concentration of 2 μM for 24 h before they were exposed to 2 or 4 Gy of γ-rays. It was shown that γ-rays alone killed breast cancer cells, and the survival rates decreased by 44.82% and 30.36% after 2 Gy irradiation to MCF-7 and MDA-MB-231 cells, respectively. After 4 Gy irradiation, the survival rates further reduced by 76.44% and 68.36% in the two cell lines (Figs 4A and B), implying that MDA-MB-231 cells were more resistant to radiation. Bosutinib alone decreased survival rates of the two cell lines by ~40%. As compared to irradiation or bosutinib alone, treatment with irradiation and bosutinib reduced the survivals of MCF-7 and MDA-MB-231 cells irradiated with 2 Gy from 55.18 ± 6.51% to 27.4 ± 0.74% and 69.64 ± 10.28% to 36.79 ± 2.05%, respectively and irradiated with 4 Gy from 23.56 ± 3.14% to 10.63 ± 1.18% and 31.64 ± 9.76% to 13.65 ± 0.79%, respectively. These data suggested that survival rates of both the breast cancer cells were the lowest when the combination of bosutinib and irradiation was used, indicating bosutinib as a promising radiosensitizer for its synergistic effect with irradiation. Thus, bosutinib may noticeably improve the efficiency of radiation in killing cancer cells.

**Fig 4.**
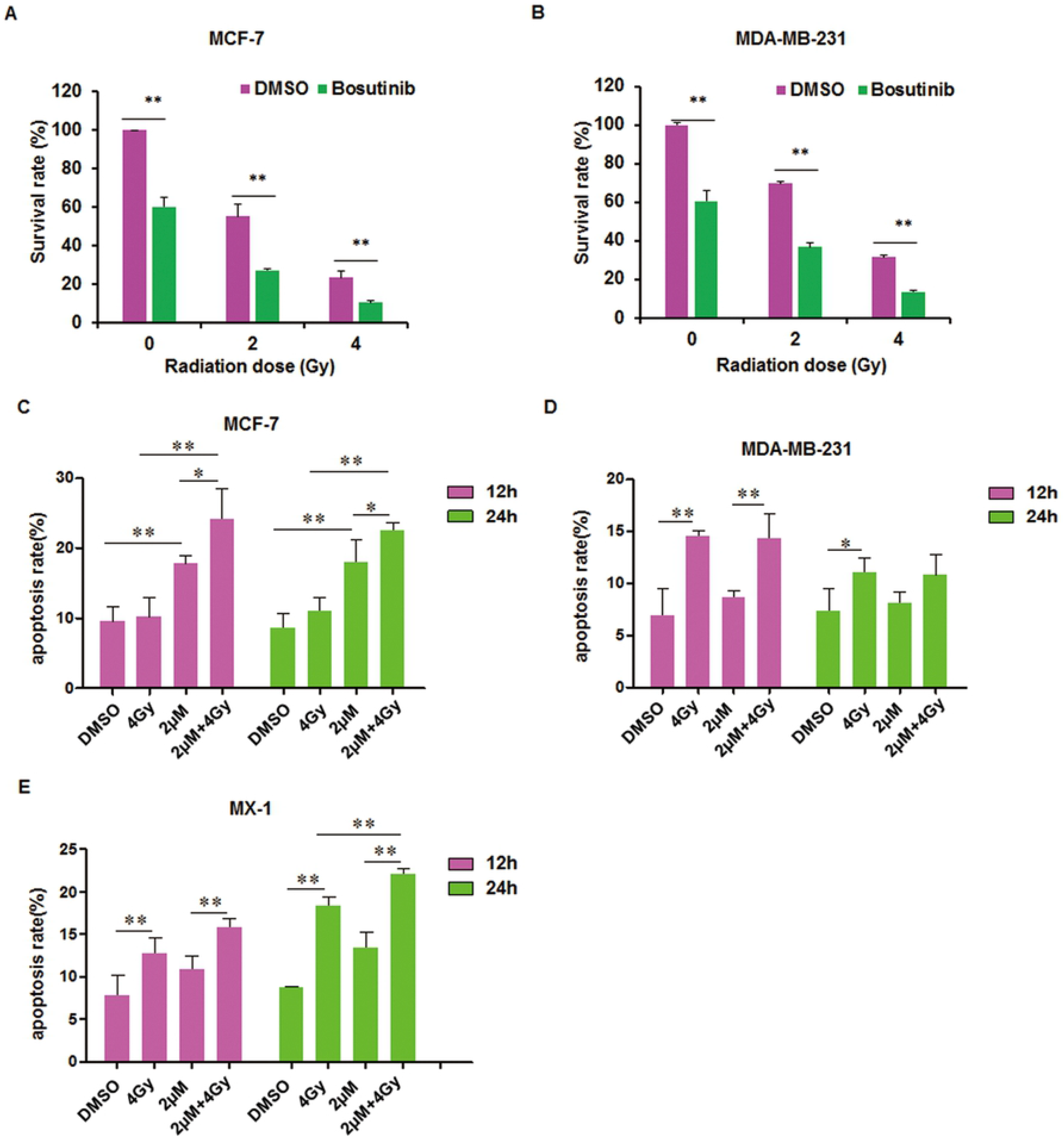
Bosutinib sensitizes cells to ionizing radiation (IR). (A-B) Sensitization of MCF-7 and MDA-MB-231 cells to γ-ray irradiation by bosutinib. (C-E) Effect of bosutinib on apoptosis induced by γ-ray irradiation in MCF-7 cells (C), MDA-MB-231 cells (D), and MX-1 cells (E). For the apoptosis assays, the cells were pre-treated with 2 μM of bosutinib for 24 h, followed by irradiation. The data were expressed as the mean and standard deviation from three independent experiments. \**P* < 0.05, \*\**P* < 0.01.

Apoptosis was detected in all three breast cancer cells at 12 h and 24 h after irradiation. After 4 Gy irradiation alone, apoptosis was significantly promoted at 12 h in MDA-MB-231 cells from 6.92% to 14.54% (*P* < 0.01, Fig 4C), and in MX-1 cells, the apoptosis increased significantly at 12 h from 7.83% to 12.77% (*P* < 0.05) and at 24 h from 8.79% to 18.40% (*P* < 0.01) (Fig 4E), but not in MCF-7 cells, indicating that MCF-7 cells were resistant to radiation (Fig 4C). The treatment with bosutinib alone induced apoptosis in MCF-7and MX-1 cells, but not in MDA-MB-231 cells. Apoptosis induced by the combined treatment of radiation and bosutinib in MCF-7 cells was 24.14% at 12 h and 22.56% at 24 h after irradiation. These apoptosis rates were much higher than that induced by radiation alone, which were nearly similar to the controls, highlighting radiosensitizing effects of bosutinib in the resistant MCF-7 cells (Fig 4D). In MX-1 cells, the addition of irradiation enhanced apoptosis from 10.96% to 15.90%, or 13.49 % to 22.15% compared with bosutinib alone at 12 h and 24 h after irradiation, respectively, further indicating the synergistic effect of bosutinib and irradiation (Fig 4E). However, bosutinib exhibited no further increased apoptosis induced by irradiation in MDA-MB-231 cells at both 12 h and 24 h after irradiation (Fig 4C).

### Bosutinib inhibits expression of *eIF4G1*

Western blot assays were conducted to detect the expression level of *eIF4G1* and its homolog *eIF4G2* in all the breast cancer cells. Fig 5A revealed that expression of *eIF4G1* in MDA-MB-231 was lower than that in the other two cell lines. In MX-1 cells, the *eIF4G1* expression level declined after treatment with bosutinib in a dose-dependent manner (Figs 5B and C). The decrease was significant at 0.05 μM and 5 μM of bosutinib (Fig 5C). However, the *eIF4G2* expression level did not change noticeably. In MCF-7 cells, the *eIF4G1* expression was significantly decreased at concentrations of more than 1 μM (Figs 5D and E). In MDA-MB-231 cells, the *eIF4G1* expression decreased significantly only at 8 μM of bosutinib (Figs 5F and G). However, bosutinib did not seem to affect the expression of *eIF4G2* in MCF-7 and MDA-MB-231 cells.

**Fig 5.**
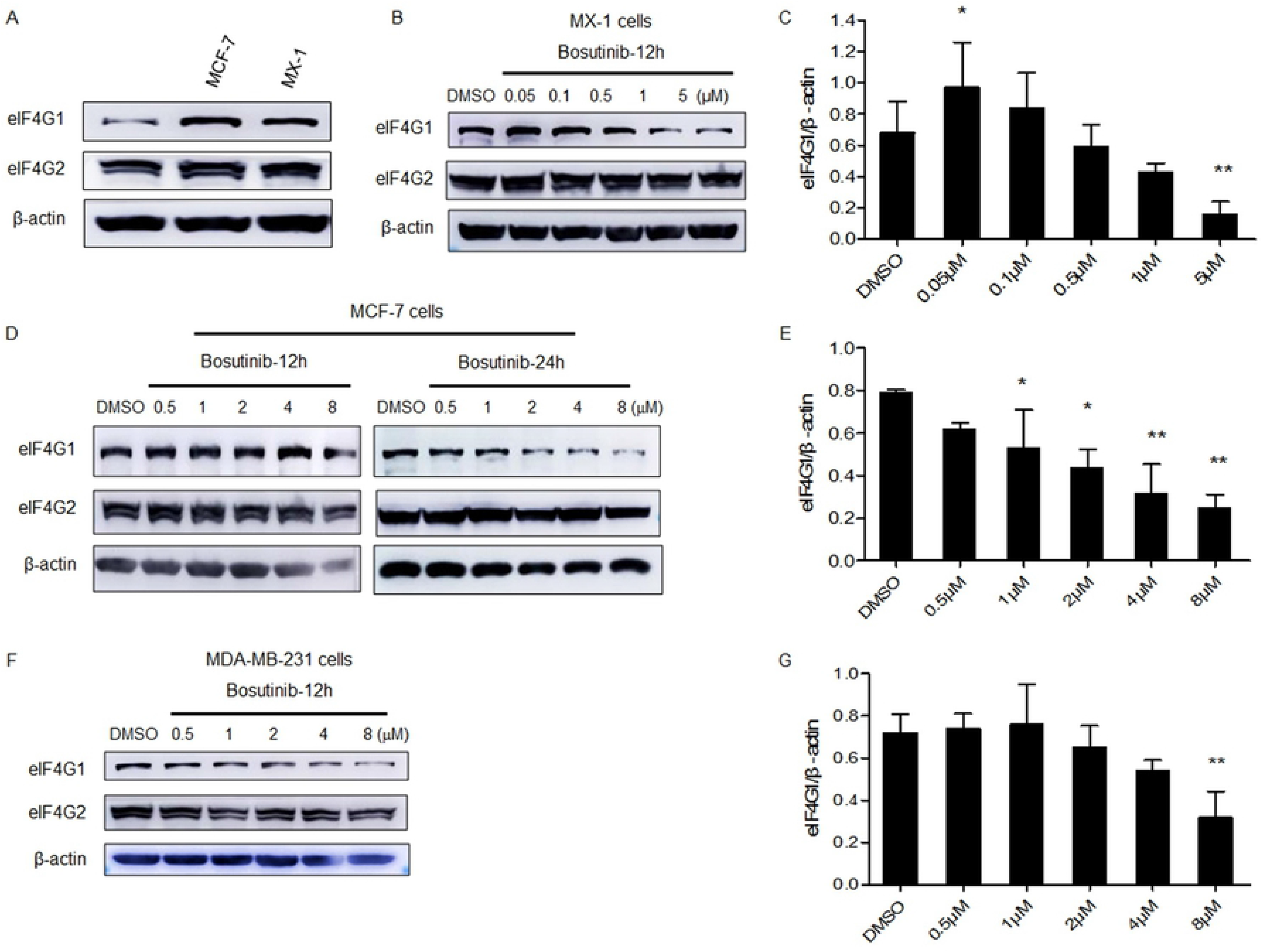
Effects of bosutinib on the expression of *eIF4G1* in different cell lines. (A) Western blot analyses of *eIF4G1* expression in the three breast cancer cell lines. (B) The alterations of *eIF4G1* expression in MX-1 cells after 12 h treatment with bosutinib at different concentrations. (C) Densitometry analyses of *eIF4G1* expression in MX-1 cells treated with bosutinib for 12 h. (D) The alterations of *eIF4G1* expression in MCF-7 cells after 12 h or 24 h treatment with bosutinib at different concentrations. (E) Densitometry analyses of *eIF4G1* expression in MCF-7 cells treated with bosutinib for 24 h. (F) The alterations of *eIF4G1* expression in MDA-MB-231 cells after 12 h treatment with bosutinib at different concentrations. (G) Densitometry analyses of *eIF4G1* expression in MDA-MB-231 cells treated with bosutinib for 12 h. The data were expressed as the mean and standard deviation from three independent experiments. \**P* < 0.05, \*\**P* < 0.01.

We further investigated the effects of bosutinib on the expression of a series of DNA damage response (DDR) proteins. We found that bosutinib increased the expression level of *γH2AX*, a biomarker of DNA double-strand break, or prolonged its vanishing effect (Fig 6A), indicating increased DNA damage induced by this drug. We further analyzed the expression changes of some other IR-induced DDR proteins. It was revealed that the expression levels of *ATM* after treatment with bosutinib were lower than before treatment at all-time points in all the three cell lines (Figs 6A, C and E). The expression levels of *XRCC4*, *ATRIP*, and *GADD45a* decreased in MDA-MB-231 and MCF-7 cells after bosutinib treatment with or without irradiation (Figs 6B and D). We also analyzed the expression of *PARP-1*, *Mre11*, *CDK1*, and *pCDK1* and found that bosutinib did not elicit apparent alteration on the expression of these proteins.

**Fig 6.**
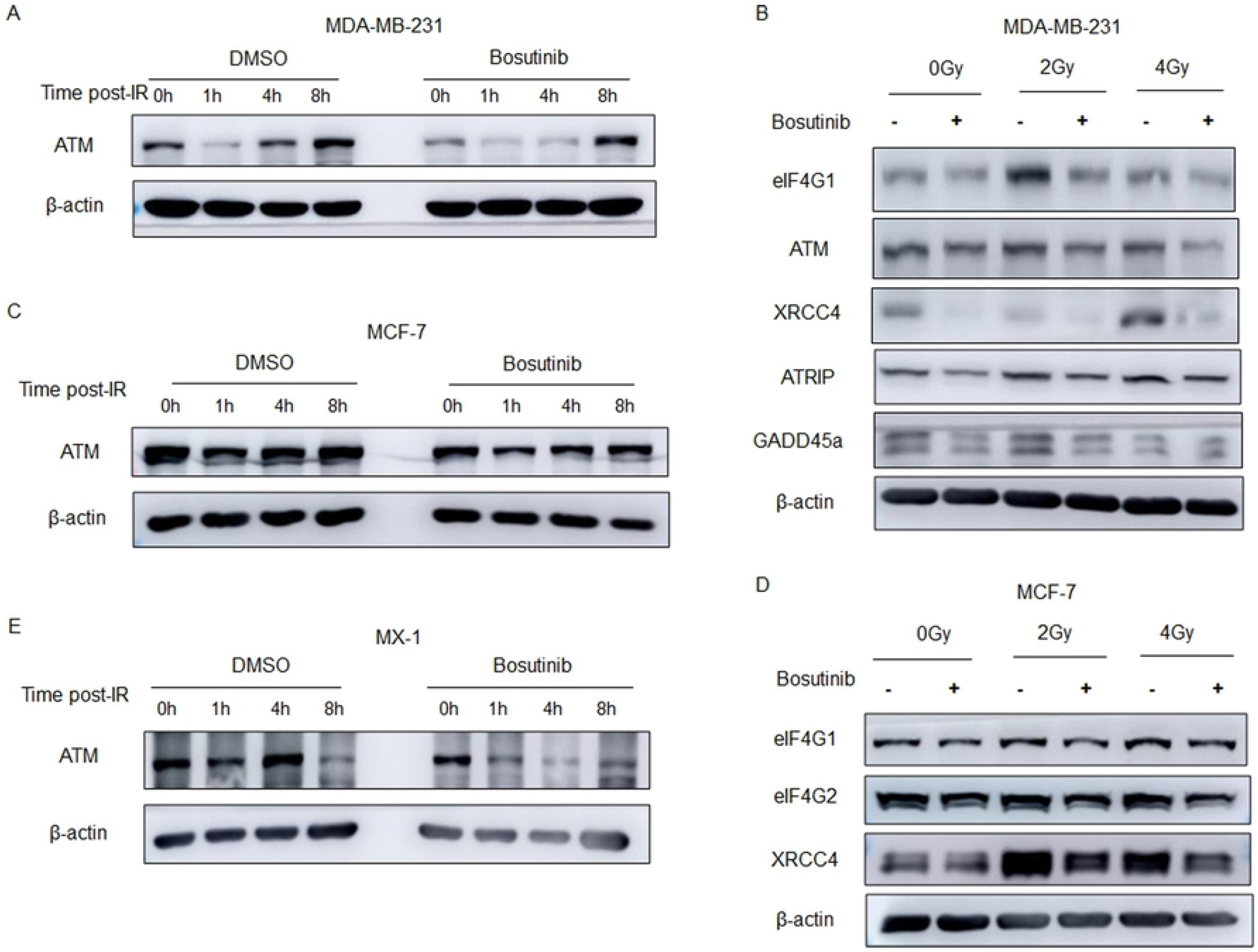
Effects of bosutinib on the expression of DNA damage response (DDR) proteins. (A) *ATM* and *γH2AX* expression in MDA-MB-231 cells. (B) Expression of DDR proteins in MDA-MB-231 cells at 12 h after irradiation. (C) *ATM* expression in MCF-7 cells. (D) Expression of DDR proteins in MCF-7 cells at 12 h after irradiation. (E) *ATM* expression in MX-1 cells. The cells were pre-treated with 2 μM of bosutinib for 24 h, then irradiated with γ-ray. Expression of DDR proteins were analyzed by Western blot assays.

## Discussion

Advances in new physical and biological techniques in RT, including radiosensitization, are endeavouring to improve the clinical effect against cancer invasion and metastasis at a relatively low dose with fewer adverse effects[27]. Therefore, effective radiosensitizers with a more favourable toxicity profile are urgently needed, and investigation into the accurate underlying mechanism of radioresistance and radiosensitization is highly desired.

*EIF4G1*, the most abundant member of the *eIF4G* family, is a large scaffolding protein in *eIF4F* complex, upon which ribosomes and eIFs assemble. Increased expression of *eIF4G1* has been frequently found to be associated with progression and metastasis in a variety of cancer types [28,29]. *EIF4G1* has been identified to possess the characteristics of a potential anticancer target.

Using computational methods, the present study investigated the repositioning opportunities of bosutinib, an oral first-line second-generation therapeutic drug for the treatment of newly diagnosed chronic myelogenous leukaemia [30], as a new sensitizer for RT and compared its gene expression signatures with that of *eIF4G1*-silenced breast cancer cells for improving the sensitivity of the breast cancer cells to irradiation. We further confirmed the regulatory role of bosutinib in radiosensitivity of breast cancer cells and primarily explored the underlying mechanism of *eIF4G1* in repairing the DNA double-strand break induced by IR. The results indicated that MCF-7 cells were highly resistant to RT in terms of induction of apoptosis compared to the other two cell lines. The order of sensitivity to bosutinib among the three breast cancer cell lines was MX-1 > MDA-MB-231 > MCF-7 cells. Apoptosis was not evidently observed in bosutinib treated MDA-MB-231 cells with or without irradiation; thus, the cell-killing could be attributed to other mechanisms of cell death such as autophagy. However, the combination treatment of radiation and bosutinib significantly enhanced the potency of cell killing. Their synergistic effect efficiently functioned at relatively low doses of both radiation and bosutinib, resulting in lower toxicity on normal tissues.

Furthermore, our results also indicated that bosutinib inhibited the *eIF4G1* expression in all three cell lines in a dose-dependent manner. Importantly, bosutinib also inhibited the expression of a series of DDR genes encoding proteins by suppressing *eIF4G1*. It was explained as the major cause of radiosensitization by targeting *eIF4G1*[20]. Consequently, the efficiency of DNA damage repair was reduced, which was displayed by the prolonged quenching effect of *γH2AX*. The mechanism of radiosensitization of bosutinib could primarily be associated with *eIF4G1*. Thus, highlighting the effectiveness of LINCS gene expression signature strategy for exploring novel applications of old drugs.

Recently, a series of studies revealed that cancer patients who had undergone RT might have increased risk of non-cancer diseases as radiation late effects, such as cardiovascular diseases[31] and stroke[32]. A comprehensive analysis demonstrated that, compared to other new-generation tyrosine kinase inhibitors, incidences of vascular and cardiac treatment-emergent adverse events in patients receiving bosutinib were markedly lower, even after long-term treatment[33]. Therefore, the combination of RT and bosutinib significantly reduces the risk of cardiotoxicity. Moreover, oral mucosa and salivary glands are very sensitive to radiation damage. Up to 40% of patients undergoing RT develop xerostomia as a result of collateral damage to salivary glands by radiation[34]. However, it has been reported that bosutinib could afford radioprotective function to the salivary gland, by blocking the activation of the pro-apoptotic action of *PKCδ* on normal tissues in patients receiving RT[35]. Therefore, the radioprotective effect of bosutinib on normal tissues may become its unique superiority as a radiosensitizer in RT.

Overall, this study describes a potent approach for identifying radiosensitizer candidates based on rational computational drug repositioning. Our findings on the regulatory mechanism of bosutinib-targeting *eIF4G1* in DDR have further expanded our understanding of gene regulation in IR-induced damage to cells. However, further animal experiments are needed to understand better the precise mechanisms overcoming the radioresistance of breast cancer. Advances in both development of effective radiosensitizers and investigation into the mechanism underlying regulatory roles of genes in IR-induced damages will improve clinical outcomes of RT and will allow enhanced therapeutic implications for overcoming radioresistance in cancer therapy.

## Conclusions

As RT plays an essential role in curative management of all stages of breast cancers, effective drugs for overcoming the evolving issue of tumor radioresistance in breast cancer therapy are urgently needed. Using computational methods, the present study proposed bosutinib, an FDA approved drug, as a potential candidate radiosensitizer for breast cancer therapy. We also have provided experimental evidences to confirm the results and revealed that the downregulation of the DDR gene set through inhibition of the *eIF4G1* expression was crucial in enhancing radiosensitivity in the breast cancer cells. Thus, bosutinib represents a potential candidate radiosensitizer for breast cancer therapy.

## Author Contributions

**Conceptualization:** PKZ, DFX, HG.

**Data curation:** XYY, DFX.

**Formal analysis:** SH, XDL, HG.

**Funding acquisition:** PKZ, DFX.

**Investigation:** SH, PKZ, DFX, BH, HG.

**Methodology:** PKZ, BH, HG.

**Project administration:** PKZ, XDL.

**Resources:** XYY, DFX.

**Software:** XYY, DFX.

**Supervision:** PKZ, HG.

**Validation:** SH, XDL, HG.

**Visualization:** PKZ, BH, HG.

**Writing - original draft:** PKZ, DFX.

**Writing - review& editing:** PKZ, DFX.

## Supporting information

**S1 Table. The maximum and average enrichment scores(ES) of known compounds compared with irradiated *eIF4G1*-silenced cells.s**

